# Analysis of Polycomb Repressive Complexes binding dynamics during limb development reveals the prevalence of PRC2-independent PRC1 occupancy

**DOI:** 10.1101/2020.09.21.306688

**Authors:** Claudia Gentile, Alexandre Mayran, Fanny Guerard-Millet, Marie Kmita

## Abstract

The Polycomb group (PcG) proteins are key players in the regulation of tissue-specific gene expression through their known ability to epigenetically silence developmental genes. The PcG proteins form two multicomponent complexes, Polycomb Repressive Complex 1 and 2 (PRC1 and PRC2), whereby the hierarchical model of recruitment postulates that PRC2 triggers the trimethylation of Histone H3 lysine 27 (H3K27me3) leading to the recruitment of PRC1. Here we report on the genome-wide binding dynamics of components from both PRC1 and PRC2 in the developing limb. We show that a large proportion of PRC-bound promoters are occupied exclusively by PRC1, suggesting a more extensive PRC1-specific activity than anticipated. We found that PRC1 (RING1B) and PRC2 (SUZ12) co-occupy the promoters of developmental genes, for which a subset become up-regulated upon the inactivation of PRC2. Strikingly, we found that RING1B occupancy is largely unaffected by the loss of PRC2, revealing a complex functional relationship between these two complexes in regulating gene expression and possibly an expansive functional interplay between canonical and non-canonical PRC1.

## Introduction

Polycomb group (PcG) proteins are known for their contribution to the silencing of developmental genes and thereby regulating cell-fate decisions. They were initially discovered in *Drosophila* for their role in repressing the *Hox* genes, which in turn regulate proper body segmentation during embryonic development. The PcG proteins form two functionally distinct complexes, Polycomb Repressive Complex 1 and 2 (PRC1 and PRC2, respectively), each known for their different catalytic activities. The PRC1 complex contains Ring1A/B which has an E3 ligase activity to catalyze the monoubiquitination of lysine 119 on histone H2A (H2AK119ub1), whereas the PRC2 complex contains Ezh2 which confers a methyltransferase activity responsible for the trimethylation of lysine 27 on histone H3 (H3K27me3). In vertebrates, the core components of PRC2 consist of Enhancer of Zeste Homologue 2 (Ezh2), Embryonic Ectoderm Development (Eed), and Suppressor of Zeste 12 (SUZ12). The PRC1 complex on the other hand, is characterized by extensive protein diversity and can be sub-classified into canonical and non-canonical PRC1 complex (cPRC1 and ncPRC1, respectively). The core of both cPRC1 and ncPRC1 complexes consists of Ring1A/B and one of the Pcgf proteins (Pcgf1-6). Additionally, only cPRC1 contains a Cbx protein (Cbx 1-8), which allows the hierarchical recruitment of cPRC1 to its targets, as the Cbx proteins recognize the H3K27me3 repressive mark deposited by PRC2 [1].

While extensive work has focused on studying the role of PcG proteins in gene silencing to prevent expression in inappropriate cells or at improper developmental times, recent evidence also points to a role of PcG proteins in gene activation [2-4]. These unexpected findings revealed that PRC1 complex composition varied in a tissue- and cell-specific manner which modifies the function of the PRC1 complex to promote gene activation [3,4]. Additionally, it has been demonstrated that the non-canonical form of PRC1 is often implicated in gene activation as it is found to occupy active genes in various developmental and pathogenic contexts [5-7]. Furthermore, the function of PRC2 in limb development is not restricted to gene silencing, as exemplified with the role of *Eed* in favoring enhancer-promoter contacts required for *HoxA* gene expression [8]. Together these findings highlight the increasing complexity behind the functional roles of the Polycomb repressive complexes in gene regulation.

The use of genome-wide approaches in embryonic stem cells to map the binding of components from each Polycomb repressive complex led to the identification of genes co-occupied by both PRC1 and PRC2, corresponding to developmental transcription factors [9,10]. However, analyses of PRC1 and PRC2 binding dynamics, *in vivo*, during developmental processes are still lacking. The developing limb is a valuable model system to investigate mechanisms of gene regulation, notably due to its accessibility for manipulation and the possibility to generate limb-specific gene inactivation without impinging on embryonic survival. The function of PRC1 and PRC2 was shown to be required for proper morphogenesis of the limb along the proximal (presumptive arm) to distal (presumptive hand) axis as well as along the anterior (thumb) to posterior axis (fifth finger) [8,11-13].

In this work, we analyzed the relationship between PRC1 and PRC2 (PRC) occupancy at promoters, genome-wide, in proximal and distal limb buds. Consistent with the role of PRC in gene silencing, a subset of PRC1 and PRC2 co-occupied promoters become transcriptionally up-regulated upon the conditional inactivation in limbs of the PRC2 core component, Eed. Our analysis also revealed that a large majority of RING1B occupancy in wild type limbs is associated with transcriptional activity, likely representative of ncPRC1 occupancy. While the inactivation of Eed does not affect the overall proximal-distal transcriptional limb identity, proper *Hox* gene expression, crucial for limb patterning, is most impacted. Finally, we uncover that RING1B occupancy is largely unaffected by *Eed* inactivation suggesting that PRC1 can be recruited or maintained in absence of PRC2.

## Results

### PRC1 and PRC2 binding dynamics in the limb establishes distinct promoter categories genome-wide

To study the binding dynamics of PRC1 and PRC2 (PRC) during limb development, ChIP-seq experiments for RING1B (PRC1) and SUZ12 (PRC2) were performed in proximal and distal limb buds at embryonic day 12.5 (E12.5) (Fig. 1A). Genome-wide ChIP-seq analysis of PRC signal intensities around transcriptional start sites (TSSs +/- 1.5kb, referred to as promoters hereafter) in proximal and distal limbs, lead to the observation that more than half of promoters are bound by PRC1 and/or PRC2 (referred to as PRC-bound promoters), with 59% in proximal limb and 56% in distal limb (Fig. 1B). Further analysis revealed a partitioning of these PRC-bound promoters into 3 distinct categories. The first category comprises promoters occupied by SUZ12 and RING1B, category 2 consists of promoters occupied by SUZ12 alone while promoters occupied by RING1B alone form category 3 (Fig. 1B). The proportions of promoters making up each category are similar in the proximal and distal limb and quite surprisingly, the largest proportion of PRC-bound promoters (79%) are exclusively occupied by PRC1 (category 3; Fig. 1C).

**Figure 1.**
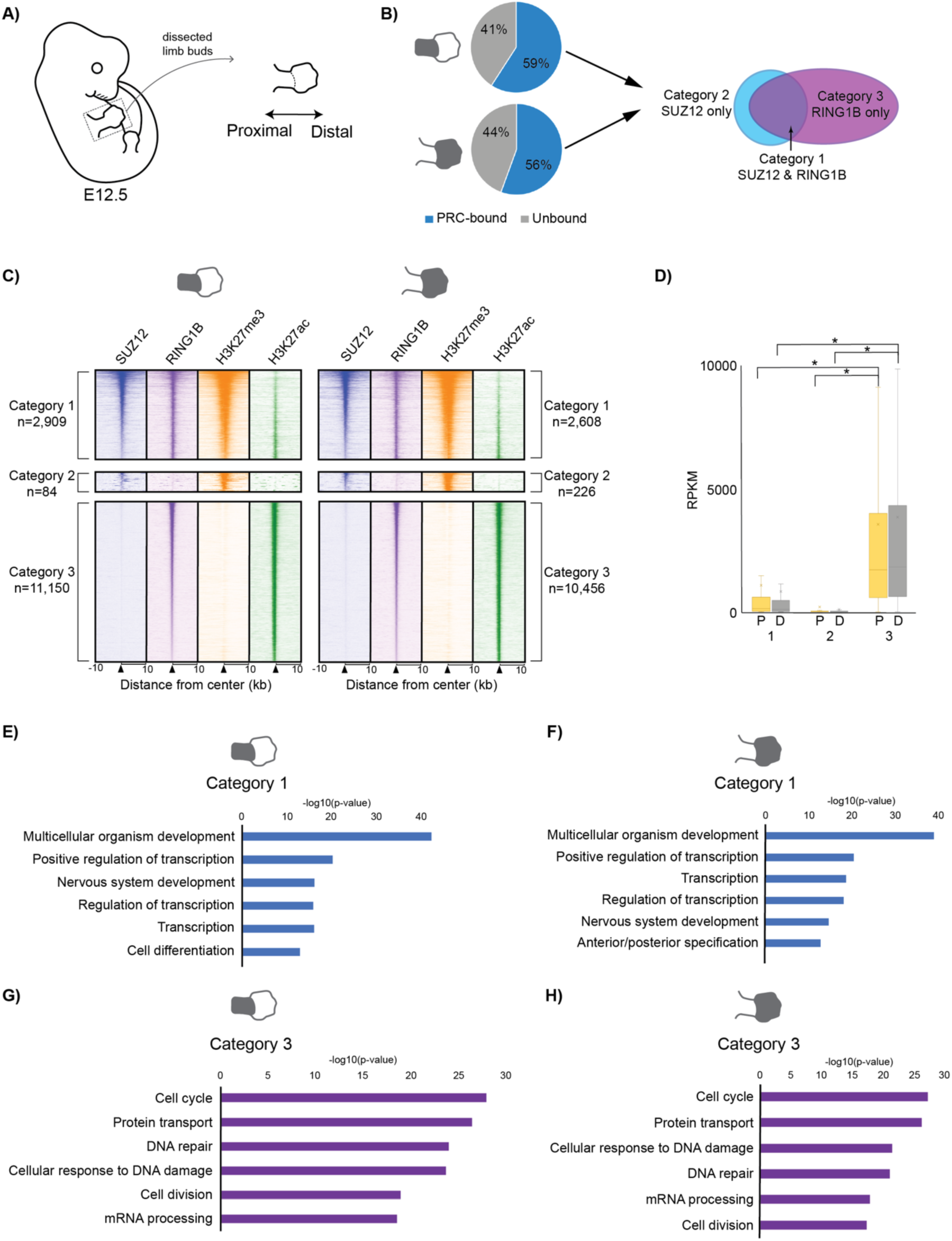
Genome-wide analysis of PRC occupancy reveals 3 distinct promoter categories. **(A)**Drawing of an embryo at embryonic day 12.5 (E12.5) used for dissection of limb buds into proximal and distal tissues **(B)** Pie charts displaying the proportion of PRC-bound vs. unbound promoters genome-wide in proximal (upper) and distal (lower) limb tissues. Arrows from PRC-bound promoters point to a diagram representing how these promoters are further broken down into 3 categories according to PRC1 and PRC2 occupancy. To note, the grey shaded areas of the limb buds represent the corresponding tissues, that is either proximal or distal. **(C)** Heatmaps of ChIP-seq data depicting the signal intensity of SUZ12 (blue), RING1B (purple), H3K27me3 (orange), and H3K27ac (green) at promoters in the proximal and distal limb for the 3 categories of promoters described. The number (n) of promoters for each category are indicated. **(D)** Box plot of gene expression (using RPKM values) from RNA-seq analysis in the proximal (P) and distal (D) limbs of each category of promoters (1-3). Significant differences (p-value <0.05) in gene expression are depicted by an asterisk and were calculated using an unpaired t-test. **(E-H)** Gene Ontology terms (using DAVID) of top 6 biological processes associated with genes from categories 1 (blue bar graphs) and 3 (purple bar graphs) in the proximal and distal limb.

SUZ12 and RING1B occupancy at category 1 promoters correlates with the presence of the H3K27me3 repressive histone mark (Fig. 1C). Importantly, within this category, H3K27me3 as well as H3K27ac signals are detected at a subset of promoters, which reflects the cell heterogeneity of the limb tissues as these two histone marks are mutually exclusive [14,15] (Fig. 1B). Category 1 promoters can thus be sub-divided into a sub-category referred to as fully-silenced promoters, that are bound by SUZ12, RING1B and are devoid of H3K27ac (i.e. 1459/2909 in proximal limb and 1525/2608 in distal limb) and a category of promoters reflecting a heterogeneous state: repressed in a subset of cells and active in the other subset (referred to as heterogeneously-silenced promoters hereafter) (Fig. S1A). The latter show ChIP-seq signals for both H3K27me3 and H3K27ac (Fig. S1A). Next, few promoters are exclusively occupied by SUZ12 in either limb tissue (category 2) and these are characterized by the presence of H3K27me3 and absence of H3K27ac (Fig. 1C). Category 3 promoters are characterized by strong RING1B and H3K27ac ChIP-seq signal intensities and appear completely devoid of H3K27me3 (Fig. 1C). As such, category 3 promoters likely correspond to promoters occupied by the non-canonical PRC1 complex (ncPRC1), which does not depend on PRC2/H3K27me3 for its recruitment to DNA [16,17]. Strikingly, the vast majority of RING1B-bound promoters belong to category 3 (approximately 80%), suggesting a high prevalence of nc-PRC1 occupancy in the developing limb.

RNA-seq analysis in wild type proximal and distal limb buds reveals that category 3 promoters coincide with highly expressed genes, in contrast to those from categories 1 and 2 (Fig. 1D). This correlates with the high H3K27ac signal intensity at category 3 promoters in comparison to category 1 and 2 promoters. Amongst the genes belonging to category 1, those showing a significant RNA-seq signal correspond to the heterogeneously-silenced promoters, which further emphasize tissue heterogeneity (Fig. S1B). Furthermore, GO analysis revealed that promoters belonging to category 1 and 3 are enriched for distinct biological processes (Fig. 1E-H) and due to the small number of promoters within category 2, GO analysis did not reveal any significant enrichment for specific biological processes. A strong enrichment for biological process terms associated with development and transcriptional regulation are found for category 1 promoters (Fig. 1E and F). This is consistent with PRC function in mediating repression of genes encoding developmental transcription factors [9,10]. In contrast, category 3 promoters, bound exclusively by RING1B, relate to genes involved predominantly in cell cycle, DNA repair and protein transport (Fig. 1G and H). These are terms that have been reported to associate with ncPRC1 targets, which has revealed the contribution of PRC1 in gene activation [6]. Overall, we show that RING1B and SUZ12 occupancy occurs at developmentally regulated genes that are silent (either in the entire limb or part of it) in comparison to the exclusive occupancy by RING1B that correlates with active promoters.

The observation that category 3 promoters, bound exclusively by RING1B, correlated so strikingly with high gene expression lead us to question whether promoters not bound by PRC (Fig. 1B) similarly correspond to highly expressed genes that in contrast do not require any PRC1 presence. Quite surprisingly, the unbound promoters demonstrated near-base line levels of gene expression when compared to category 3 gene expression (Fig. S1C-D). GO analysis for these unbound promoters represent biological process terms associated with sensory perception and signal transduction, amongst other similar top hits (Fig. S1E-F). The implications of this finding are two-fold: firstly, these genes do not require active silencing by PRC binding at their promoters to be in this “repressed state” (at least in this cell type) and secondly, the most transcriptionally active genes genome-wide are associated with RING1B binding at their promoters.

### Transcriptional upregulation following *Eed* inactivation results from a gain of activity at nearby enhancers

To assess the genome-wide functional impact of PRC occupancy on transcriptional activity, we studied the effects of *Eed* inactivation using the previously validated *Eed* limb-conditional mutant mouse line (*PrxCre; EedFlox/-*, referred to as *Eed*^*c/-*^) [8]. Importantly, the inactivation of *Eed* leads to a significant genome-wide reduction in H3K27me3 levels in both proximal and distal limb tissues (Figure S2A). Analyzing changes in transcriptional activity upon *Eed* inactivation on a genome-wide scale revealed that the majority of genes (93% and 86% in proximal and distal tissues, respectively) are unaffected by the loss of PRC2 function (Fig. S2B). This result is consistent with the observation that the vast majority of PRC-bound promoters are bound exclusively by RING1B and are in an active state in the wild type context (Fig. 1C and 1D). We therefore focused the analysis on the effect of *Eed* inactivation specifically on category 1 genes (i.e. bound by PRC1 and PRC2) in both the proximal and distal limb. Comparison of RNA-seq data in wild type and *Eed*^*c/-*^ limbs demonstrates that a large proportion of category 1 genes are also unaffected (71% and 58% in proximal and distal limbs, respectively), only a subset become significantly up-regulated (24% in proximal limb and 39% in distal limb) and a considerably smaller subset become down-regulated (5% in proximal limb and 3% in distal limb) (Fig. 2A). Significantly up-regulated genes correlate with an increase in H3K27ac histone mark at their promoters (Fig. 2B-C). To note, the up-regulated genes correspond to genes from both subsets of category 1 promoters, either fully-silenced or heterogeneously-silenced genes in the wild type limb, with a similar proportion of genes becoming up-regulated in either of the two sub-categories (Fig. S2C). In agreement with these findings, our analysis indicates that PRC2 function is required to prevent the up-regulation of genes that are either expressed in a subset of proximal and/or distal cells (heterogeneously-silenced), such as *Meis1*, or of genes that are entirely silent in the proximal and/or distal cells (fully-silenced), such as *Hoxb13, Hoxd13* and *Hoxc11* (Fig. 2D-E). These results are consistent with PRC2 function being required for ensuring gene silencing in a spatially-regulated manner. However, a large proportion of genes within category 1 are unaffected by *Eed* inactivation in the proximal and distal limb, which suggests that loss of PRC2 binding at gene promoters is not sufficient for their activation.

**Figure 2.**
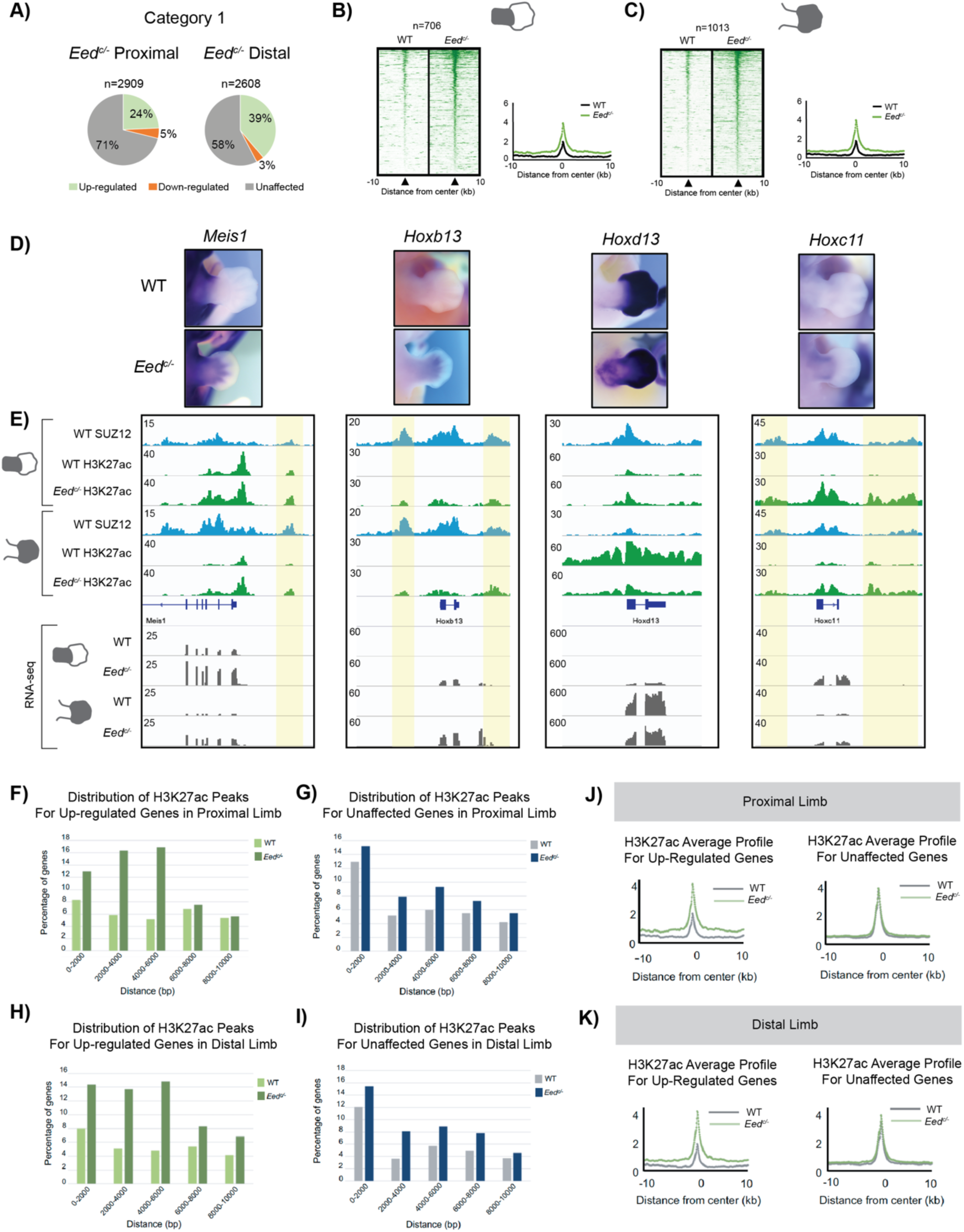
Transcriptional up-regulation results from a gain in enhancer activity upon PRC2 inactivation in the limb. **(A)** Pie charts representing the proportions of genes bound by PRC1 and PRC2 at their promoters that are unaffected, up-regulated (FDR<0.05, FC>0) or down-regulated (FDR<0.05, FC<0) upon PRC2 inactivation in the proximal limb and distal limb. **(B-C)** Heatmaps of H3K27ac ChIP-seq at PRC1/2 bound targets that are significantly (FDR<0.05) up-regulated in *Eed*^*c/-*^ proximal (B) and *Eed*^*c/-*^ distal limbs (C) with corresponding average profiles. **(D)** Whole-mount *in situ* hybridizations of *Meis1, Hoxb13, Hoxd13* and *Hoxc11* expression in wild type and *Eed*^*c/-*^ limbs. **(E)** IGV snapshots of SUZ12 and H3K27ac ChIP-seq tracks and RNA-seq at up-regulated genes in wild type and *Eed*^*c/-*^ proximal and distal limbs. Gains in H3K27ac at putative enhancers are depicted by the yellow shadowbox. **(F-I)** Bar graphs representing the distance of H3K27ac peaks to promoters of genes that are either up-regulated or unaffected by PRC2 inactivation in proximal (F, G) and distal (H, I) limbs. **(J, K)** Average profiles of wild type and *Eed*^*c/-*^ H3K27ac for up-regulated and unaffected genes in proximal (J) and distal (K) limbs.

We observe that the gain of H3K27ac at up-regulated genes is also associated with a gain of H3K27ac at non-coding loci located upstream and/or downstream these genes, likely representing regulatory elements. Interestingly, these non-coding loci are bound by SUZ12 in the wild type limb (Figure 2E; yellow shadowboxes). This led us to question if genes that become up-regulated versus genes that are unaffected by *Eed* inactivation are more likely to become associated with active enhancers. To do so, for each upregulated and unaffected gene promoter, we analyzed the distance to the closest H3K27ac peak under wild type and *Eed*^*c/-*^ conditions. In wild type tissues, H3K27ac marks are at relatively similar distances from target promoters regardless of whether they become up-regulated or unaffected by *Eed* inactivation (Fig. S2D). Thus, it does not appear that pre-existing active regulatory elements influence the transcriptional response of PRC-bound genes to *Eed* inactivation that would otherwise pre-determine which genes become up-regulated. In contrast, we observe in *Eed*^*c/-*^ limb buds, that up-regulated genes are more frequently associated with H3K27ac peaks at closer distances to their promoters as compared to unaffected genes (Fig. 2F-I). In fact, we find that in both proximal and distal limbs, the percentage of up-regulated genes that are associated with H3K27ac peaks within a 6 kb window more than doubles upon *Eed* inactivation, as compared to unaffected genes (Fig. S2E). Such gain in H3K27ac in the vicinity of up-regulated genes likely reflects a gain in activity of regulatory elements. The lack of a gain in H3K27ac near genes transcriptionally unaffected by *Eed* inactivation suggests that loss of *Eed* function is not sufficient for triggering a transcriptional up-regulation of genes normally occupied by PRC2.

### PRC2 is required for proper *Hox* expression but not for proximal-distal identity during limb development

Our data demonstrates that PRC1 and PRC2 co-target genes involved in developmental processes and that a subset of these genes become misregulated upon loss of PRC2 function. Therefore, we next addressed the extent to which these transcriptional changes impact limb morphogenesis along the proximal to distal axis, since we previously showed defects along this axis upon inactivation of *Eed* [8]. To assess the contribution of PRC2 function on the transcriptional programs that characterize the proximal *versus* distal limb (i.e. the differentially expressed genes), we analyzed the transcriptome of proximal and distal *Eed*^*c/*-^ limb buds in comparison to wild type limb buds. Differentially expressed genes were determined by comparing the wild type RPKM levels in the proximal and distal limb buds and genes that were significantly (FDR <0.05) more expressed in one tissue versus the other limb tissue were assigned as being a differentially expressed gene along the proximal to distal axis. Clustering of the top 1000 most differentially expressed genes into a heatmap, generates four clusters representing the wild type and *Eed*^*c/-*^ proximal and distal limbs (Fig. 3A). Each cluster can be further divided by gene expression levels (orange vs blue). Comparison of the wild type limb tissues demonstrates that genes which are expressed, for example distally (orange cluster), are not found to be significantly expressed in the proximal limb tissue (blue cluster) and vice versa, emphasizing that these are differentially expressed gene sets (Fig. 3A). Interestingly, comparison of wild type vs *Eed*^*c/-*^ clusters within a tissue, reveals that the overall clustering of genes in the proximal and distal limb is unaffected in *Eed*^*c/-*^ limbs, with only a very small subset of genes showing transcriptional changes, as highlighted at the bottom of the heatmap (Fig. 3A, red arrow). Accordingly, Principal Component Analysis (PCA) of the top 1000 differentially expressed genes shows that the overall proximal *versus* distal limb identity is mostly found in component 1 (88% of total variance) while transcriptional changes that occur in *Eed*^*c/-*^ limbs is found in component 2 (6.5% of total variance) (Fig. 3B). This PCA, together with the heatmap analysis, establish that the most differentially expressed genes are primarily changed between proximal vs distal limb tissues rather than between wild type vs *Eed*^*c/-*^ limb tissues. In parallel, PCA of the most significantly (FDR <0.05) misregulated genes, as determined by comparison of proximal and distal *Eed*^*c/-*^ expression data this time around, similarily reveals that most variance lies along component 1 (72.7% of total variance), which represents changes between proximal vs distal limb tissue rather than between wild type vs *Eed*^*c/-*^ limb tissue (Fig. 3C). Taken together, PCA using these two approaches, reveal that proximal-distal transcriptional limb identity is, for the most part, unaffected by the inactivation of *Eed* (Fig. 3A-C).

**Figure 3.**
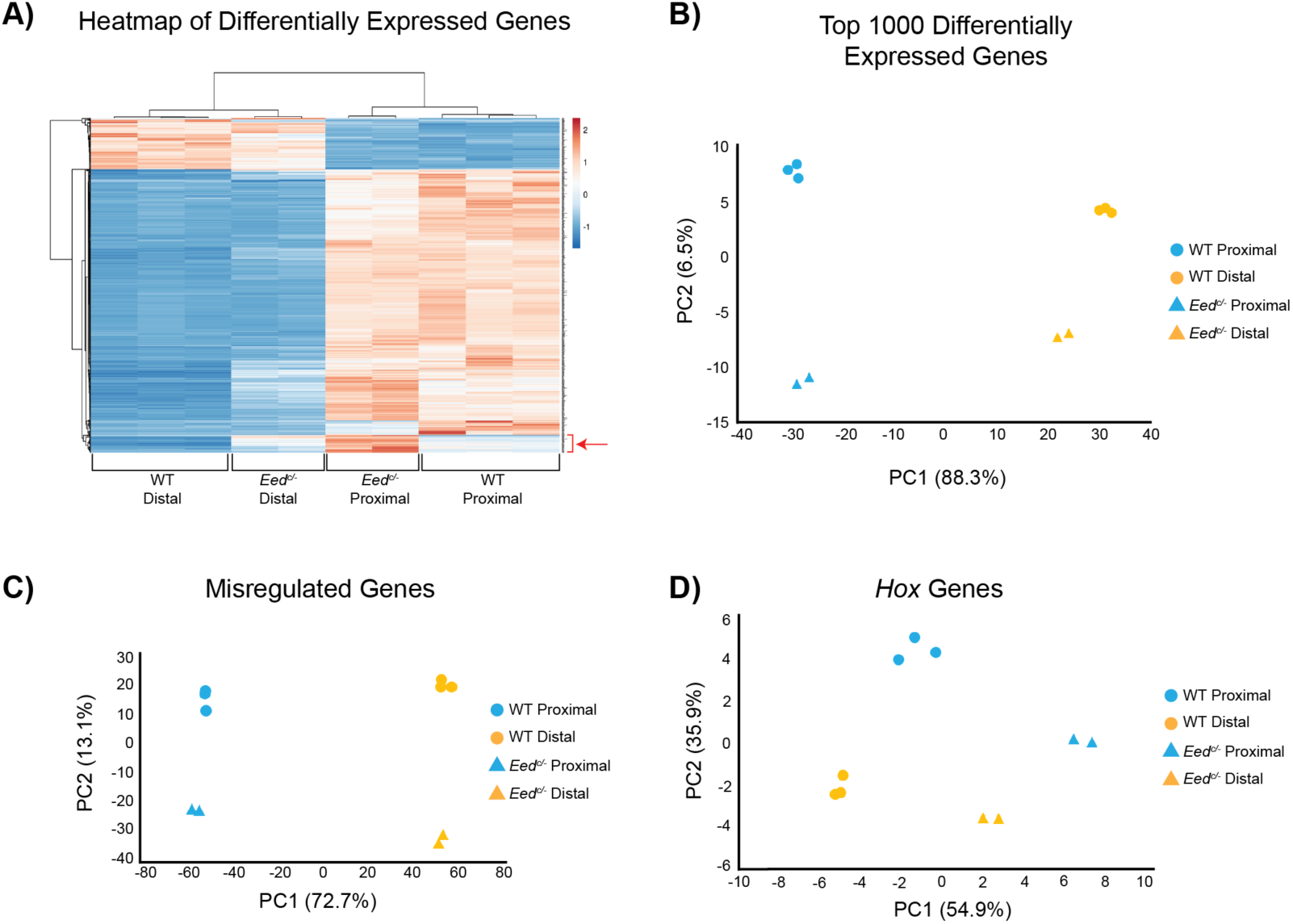
PRC2 inactivation affects *Hox* gene expression but not the overall proximal-distal limb identity. **(A**) Heatmap hierarchical clustering of top 1000 differentially expressed genes using the ClustVis tool. Clustering was performed using gene expression (RPKM) values between wild type and *Eed*^*c/-*^ proximal and distal limb tissue samples, with expression levels according to the orange-blue scale. Four clusters emerge and represent distal limb clusters on the left and proximal limb clusters on the right. At the bottom of the heat map, the red bracket and arrow point to gene expression changes occurring when comparing wild type to *Eed*^*c/-*^ limb clusters. **(B-D)** Principal Component Analysis (PCA) plots of the top 1000 differentially expressed genes (B), misregulated genes (includes up- and down- regulated genes) upon comparison of *Eed*^*c/-*^ proximal and distal limb (C), and of *Hox* gene expression (D).

Further analysis of the subset of genes that are changing in their clustering upon inactivation of *Eed*, uncovered the *Hox* genes as being amongst some of the most prevalent. This is visualized by similar PCA, which shows *Hox* genes as being mis-expressed in both the proximal and distal limb in the *Eed*^*c/-*^ mutant condition, such that more variance lies along component 1 (∼55%), which in this case characterizes the variances between the wild type and *Eed*^*c/-*^ tissues rather than between the wild type proximal and distal limb tissues. This PCA also demonstrates that *Eed*^*c/-*^ proximal limb categories cluster closer to *Eed*^*c/-*^ distal categories and away from the wild type proximal cluster, which suggests that the ‘*Hox* expression identity’ in the proximal limb is most affected by *Eed* inactivation (Fig. 3D). This analysis correlates with the impaired regulation of the *Hox9, 10* and *11* genes which are important for patterning the proximal limb (Fig. S3) [8]. Overall, our analysis of differentially expressed genes upon *Eed* inactivation in the developing limb, emphasizes the important contribution of PRC2 in regulating *Hox* gene expression to ensure proper proximal-distal limb identity.

### RING1B Occupancy Appears Unaffected by *Eed* Inactivation in the Limb

Finally, we assessed the impact of *Eed* inactivation on PRC1 recruitment by monitoring the changes in RING1B occupancy. When comparing RING1B occupancy in wild type and *Eed*^*c/-*^ limb tissue on a genome-wide scale, we observe that, quite strikingly, RING1B occupancy appears largely unchanged (Fig. 4A). When focusing on RING1B occupancy status specifically at promoters, RING1B occupancy is unaffected at category 3 promoters (devoid of PRC2 in the wild type limb tissue), as anticipated, and remain strongly associated with H3K27ac (Fig. S4). Interestingly, RING1B occupancy remains largely unaffected upon *Eed* inactivation at category 1 promoters, both in proximal and distal forelimb buds (Fig. 4B and 4C). This is a counterintuitive outcome given the proposed sequential recruitment model whereby PRC2 binding catalyzes H3K27me3, which in turn is recognized by cPRC1, resulting in PRC1 and PRC2 co-occupancy at target genes [18]. Based on the absence of the H3K27me3 mark in *Eed*^*c/-*^ limb buds (Fig. S2A), the RING1B occupancy observed indicates that it coincides with PRC1 recruitment occurring independently of the canonical mechanism. Moreover, we observe that RING1B occupancy is unambiguously detectable at genes that become up-regulated as well as at genes unaffected by *Eed* inactivation (Fig. 4B and 4D), questioning the functional role of PRC1 at these sites. Our finding of RING1B occupancy at genes that become up-regulated in *Eed*^*c/-*^ limb buds and showing a gain of H3K27ac (Fig. 4B) is consistent with the recent evidence that ncPRC1 complex can be associated with transcriptional activation [5-7].

**Figure 4.**
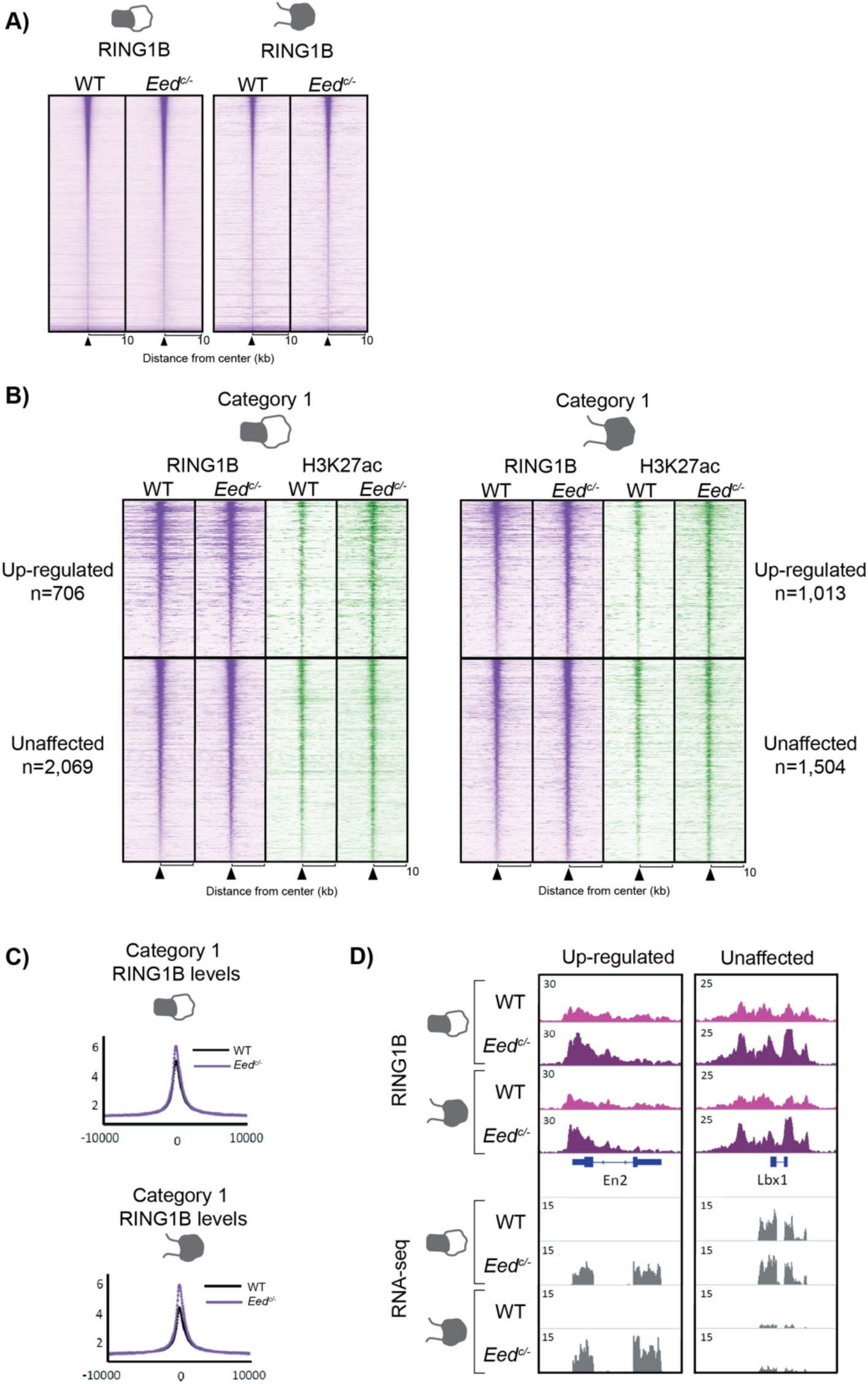
RING1B occupancy is unaffected by PRC2 inactivation. **(A)** Heatmap representation of genome-wide RING1B peaks in wild type and *Eed*^*c/-*^ proximal and distal limb. **(B)** Heatmaps of RING1B (purple) and H3K27ac (green) ChIP-seq in wild type and *Eed*^*c/-*^ proximal and distal limbs at promoters occupied by both PRC1 and PRC2 (category 1) separated according to whether genes are up-regulated or unaffected by the inactivation of *Eed*. **(C)** Average profiles comparing wild type and *Eed*^*c/-*^ RING1B levels in proximal (upper) and distal (lower) limbs. **(D)** IGV snapshots for RING1B ChIP-seq and for RNA-seq at the up-regulated *En2* gene and at the unaffected *Lbx1* gene in wild type and *Eed*^*c/-*^ proximal and distal limbs.

## Discussion

The Polycomb group proteins play important roles in gene silencing and are essential during embryonic development. As such, understanding the mechanism(s) underlying the dynamics of PRC recruitment during embryogenesis has become a major undertaking in the field. In our work, we show that PRC1 and PRC2 co-occupy a subset of promoters, primarily promoters of developmental genes, including the *Hox* gene family. The presence of both PRC1 and PRC2 at target sites has been explained by the hierarchical recruitment model, as supported by studies reporting the recognition of H3K27me3 by PRC1 [19,20]. However, numerous studies have revealed that the relationship between PRC1 and PRC2 is more complex than this model proposes, as increasing evidence indicates that PRC1 can be targeted independently of PRC2/H3K27me3 and also suggests a role for PRC1 and H2AK119ub1 in PRC2 recruitment [16,17,21,22].

Our analysis of PRC1 (RING1B) and PRC2 (SUZ12) occupancy at promoters in the developing limb revealed 3 distinct PRC occupancy patterns, 20% of which corresponds to the occupancy by both SUZ12 and RING1B at promoters (category 1). In line with this work being done in the developing limb, these sites represent largely developmental genes, amongst which we find the *Hox* genes. Polycomb group protein function at the *Hox* clusters has been extensively studied and has provided insights into the collinear regulation of *Hox* genes during developmental processes [23]. The patterning of the developing limb relies largely on the *HoxA* and *HoxD* genes [24] and their deregulation leads to limb malformations [25]. Previous studies reported the functional importance of RING1A/B proteins and EZH2 in proximal-distal limb morphogenesis and axis elongation, respectively [11-13]. Moreover, *Ezh2* inactivation in the developing limb leads to the deregulation of *Hox* genes, predominantly the paralogous group genes *9* to *13* of the *HoxA* and *HoxD* clusters, which likely contributes to the observed shortening and abnormal shape of the limb segments [11]. It was thus proposed that *Ezh2* is essential for proximal-distal morphogenesis by preventing the ectopic expression of *Hox* genes at early developmental time points, characterized by cell fate plasticity [11]. By analyzing differentially expressed proximal and distal genes, our study on *Eed* inactivation reveals that a predominant effect of PRC2 loss-of-function in the developing limb is the deregulation of *Hox* genes, particularly the *Hox9, 10* and *11* genes involved in patterning the zeugopod (forearm). While the overall proximal to distal limb segmentation into three domains remains unaffected in *Eed* mutant limbs, there are clear defects in the shape and size of the distinct limb segments, indicating that PRC2 function is required to refine gene expression in the three limb segments while segment specificity remains largely independent of PRC-dependent gene regulation. Importantly, the most drastic effect of PRC2 inactivation on *Hox* regulation is the down-regulation of *Hox9, 10* and *11* as demonstrated in this work and by others [8,11]. In fact, our results reveal that this down-regulation is not restricted to *Hox* genes. This could represent indirect effects of PRC2 inactivation in which the loss of PRC2 results in the up-regulation of a transcriptional repressor, and thereby the down-regulation of its target genes. Alternatively, the down-regulation of certain genes may be the result of alterations in genome topology. Indeed, in addition to the well-known role of PRC in mediating long-range PRC-dependent contacts involved in gene silencing [26-28], several studies have uncovered a role for PRC in organizing chromatin contacts associated with gene activation. The latter includes PRC-dependent pre-setting of enhancer-promoter contacts [29-31] and PRC-dependent 3D chromatin organization which favours physical proximity between active enhancers and promoters (not bound by PRC2) [8].

Interestingly, we demonstrate that RING1B occupancy is unaffected by PRC2 inactivation at loci co-occupied by PRC1 and PRC2 in the wild type limb. This finding was unexpected when considering the classic recruitment model for PRC1 by recognition of the H3K27me3 mark deposited by PRC2 at loci bound by both complexes. Based on the significant absence of the H3K27me3 mark in *Eed*^*c/-*^ limb buds, this RING1B occupancy most likely reflects the presence of ncPRC1, which could either result from ncPRC1 being already present at these loci in the wild type context or corresponds to *de novo* recruitment of ncPRC1 in absence of cPRC1 recruitment. In fact, ChIP-seq studies for different proteins from either canonical or non-canonical PRC1 has provided insight into the behaviour of their binding patterns whereby both cPRC1 and ncPRC1 can share the same targets [6,7,32,33] and therefore multiple complexes may operate at the same target loci. Whether distinct PRC1 complexes co-occupy the same locus or whether there is molecular heterogeneity within cell populations examined so far remains unclear. Nonetheless, these findings propose that different modes of gene regulation are at play depending on which variation of PRC1 is present at a given locus. The question now arises as to how the activity of these different complexes is coordinated to promote either gene repression or activation when all forms (canonical and non-canonical) are present at a given locus.

Furthermore, our data reveals that, in the wild type developing limb, about 80% of RING1B occupancy at promoters is independent of PRC2 (category 3). This occupancy is thus likely representative of non-canonical PRC1 as these promoters are devoid of PRC2 and H3K27me3 and are highly active and acetylated on H3K27. This is in line with similar findings reporting ncPRC1 at active genes [5-7] and points to a model where PRC1 function may be primarily related to gene activation. However, upon inactivation of PRC2, we report RING1B occupancy at genes that are both up-regulated as well as transcriptionally unaffected, questioning the role of RING1B at genes that do not become signficantly active. The prominent PRC2-independent PRC1 occupancy reported here raises the possibility that cell differentiation towards final cell fate requires PRC1-specific function. In line with these results, evidence was obtained in mESCs supporting the role of ncPRC1 in gene repression [34] even in the absence of PRC2 [35]. This may explain why the subset of genes, bound by PRC2 in the wild type context, remain transcriptionally silent upon PRC2 inactivation in the limb. Another intriguing possibility may be a lack of enhancer activation to complement the occupancy of RING1B at promoters unaffected by PRC2 inactivation, as our genome-wide H3K27ac analysis uncovers a correlation between genes that become up-regulated and a gain in enhancer activity. Nonetheless, how PRC1 can promote both gene repression and gene activation remains an intriguing question, which most likely reflects distinct functional properties depending on PRC1 subunit composition (reviewed in e.g. [36]). Further experimentation to establish the differences in genomic localization between the cPRC1 versus ncPRC1 in wild type and *Eed*^*c/-*^ limbs will likely shed light on the contribution of different complex compositions in gene regulation, but given the diversity of ncPRC1 composition, this is a challenging task.

In summary, the set of data presented here points to the predominant PRC2-independent activity of PRC1 in the developing limb, which appears largely associated with transcriptionally active genes. This is consistent with the results of Pachano et al.,[31] showing that long-range enhancer-gene interactions most likely relies on PRC1-dependent contacts between PRC-occupied orphan CpG Islands (CGIs) located near enhancers and CGIs at gene promoters. Together, these results raise the possibility that PRC1 contribution to gene activation could actually be a rule rather than an exception. Based on the selective eviction of PRC2 but persistence of PRC1 occupancy being associated with gene activation presented in this work and by Pachano et al. [31], it is tempting to speculate that the co-occupancy by PRC1 and PRC2, both at enhancers and promoters, is critical for swift transitions from silencing to transcriptional activation, notably for developmental genes.

### EXPERIMENTAL MODEL AND SUBJECT DETAILS

The inactivation of *Eed* was generated using the limb specific *PrxCre* line [37] crossed with the previously generated *Eed* floxed mouse line [38] (Jax stock #022727).

### METHOD DETAILS

#### Whole-mount in situ hybridizations

*In situ* hybridizations were carried out according to standard procedures as previously described [8]. The RNA probe for *Meis1* was as described previously [39] and the *Hoxd9-13* RNA probes were the probes described in [40]. The RNA probes for *Hoxb13* and *Hoxc11* were obtained by PCR amplification of limb cDNA using the following primers:

*Hoxb13*_F:CGAAGGGATCCGCAATTATGCCACCTTGGAC

*Hoxb13*_R:CGAAGTAATACGACTCACTATAGAAGCTTCAAACGCTGCTTTCCAGAAT

*Hoxc11*_F: AACCTCTATCTGCCCAGTTGC

*Hoxc11*_R: TAATACGACTCACTATAGGGCACTTGTCGGTCTGTCAGGTT

#### RNA Preparation and Sequencing

RNA was extracted from dissected proximal and distal limb buds from 3 independent pools of wild type limb buds (n=3) and from two independent pools of *Eed*^*c/-*^ limb buds (n=2). The extraction was performed using the RNAeasy Plus mini kit. Next, mRNA enrichment, library preparation and flow-cell preparation for sequencing were performed by the IRCM Molecular Biology Core Facility according to Illumina’s recommendations. Sequencing was done on a HiSeq 2000 instrument with a paired-end 50 cycles protocol.

#### Chromatin Immunoprecipitation (ChIP) and Sequencing

ChIP-seq experiments for SUZ12, RING1B, and H3K27ac were performed as previously described [8]. ChIP-seq experiments for H3K27me3 in proximal and distal wild type and *Eed*^*c/-*^ limb tissue were performed by crosslinking tissue with 1% formaldehyde for 13 minutes and sonicated to obtain fragments between 100-600 bp. Protein A and Protein G Dynabeads (Invitrogen) were incubated for 6 hours at 4°C with 5 μg H3K27me3 (07-449, Millipore) and coupled to beads overnight at 4°C. The samples were washed 3 times using the following buffer: 100mM Tris (pH 8), 500mM LiCl, 1% NP-40, 1% Na-Deoxycholate. The DNA was then purified on QIAquick columns (Qiagen) and sequencing was performed on an Illumina HiSeq 2000 sequencer in a 50 cycles paired-end configuration.

### QUANTIFICATION AND STATISTICAL ANALYSIS

#### ChIP-seq Data Analysis

ChIP-seq reads were aligned to the mm10 genome using bowtie v.2.3.1 with the following settings: bowtie2 -p 8 --fr --no-mixed --no-unal –x. Sam files were converted into tag directories using HOMER v4.9.1 and into bam files using Samtools v1.4.1 view function. Analysis of genome-wide occupancy for H3K27ac, H3K27me3 and RING1B was done by identifying peaks for these datasets by comparing each sample to its input using MACS v2.1.1.20160309 callpeak function using the parameters: --bw 250 -g mm --mfold 10 30 -p 1e-5. Analysis of SUZ12, H3K27me3, RING1B and H3K27ac promoter occupancy was determined using signal intensities and by averaging values from ChIP-seq data replicates (for distal limb datasets of SUZ12 and RING1B only). Heatmaps and average profiles were generated using the Easeq software (http://easeq.net) [41]. ChIP-seq data were visualized on the IGV software using BigWig files generated using the makeUCSCfile HOMER command using the following parameters: -fsize 1e20 -fragLength 150 –bigWig. GO analysis was performed using the DAVID webtool [42,43].

#### RNA-seq Data Analysis

RNA-seq experiments were carried out as previously described [8]. Paired-end reads were aligned to the mm10 reference genome using TopHat2 [44] with the parameters --rg-library “L” --rg-platform “ILLUMINA” --rg-platform-unit “X” --rg-id “run#many” --no-novel-juncs --library-type fr-firststrand -p 12. Gene expression was quantified with the HOMER analyzeRepeats command [45] and differential expression was assessed with edgeR 3.12.1[46]. MA plots were generated by averaging the wild type and *Eed*^*c/-*^ counts per million (CPM) and plotting the log(CPM+1) against the log2 fold-change (log2FC) of wild type vs *Eed*^*c/-*^ CPM values. The heatmap clustering and PCA was performed using the ClustVis webtool [47].

### DATA AND SOFTWARE AVAILABILITY

The ChIP-seq data sets used for analysis of SUZ12, RING1B and H3K27ac occupancy in proximal and distal limb tissues at E12.5 have been deposited in GEO under the accession number GSE123630.

## Acknowledgements

We thank Odile Neyret and members of the molecular biology and functional genomics core facility at the IRCM for generating sequencing libraries. We are particularly grateful to Kmita lab members for helpful discussions. Bioinformatics analyses were enabled in part by support provided by Calcul Quebec (www.calculquebec.ca) and Compute Canada (www.computecanada.ca). This work was supported by the Canadian Institute for Health Research (MOP-115127) to MK. CG was supported by the Jacques Gauthier IRCM fellowship, AM was supported by the IRCM Challenge Fellowship and FGM is supported by the IRCM Challenge Fellowship.

## Author Contributions

CG and MK conceived the study. CG designed and performed the experiments. FGM helped with limb bud dissections, performed ChIP-seq experiments for H3K27me3 and performed in situ hybridizations for *Hoxc11*. AM performed the RNA-seq analysis. CG, AM and MK analyzed and interpreted the data. CG and MK wrote the manuscript with input from AM.

