## Supplemental Figures for "Analysis of Polycomb Repressive Complexes binding dynamics during limb development reveals the prevalence of PRC2-independent PRC1 occupancy"

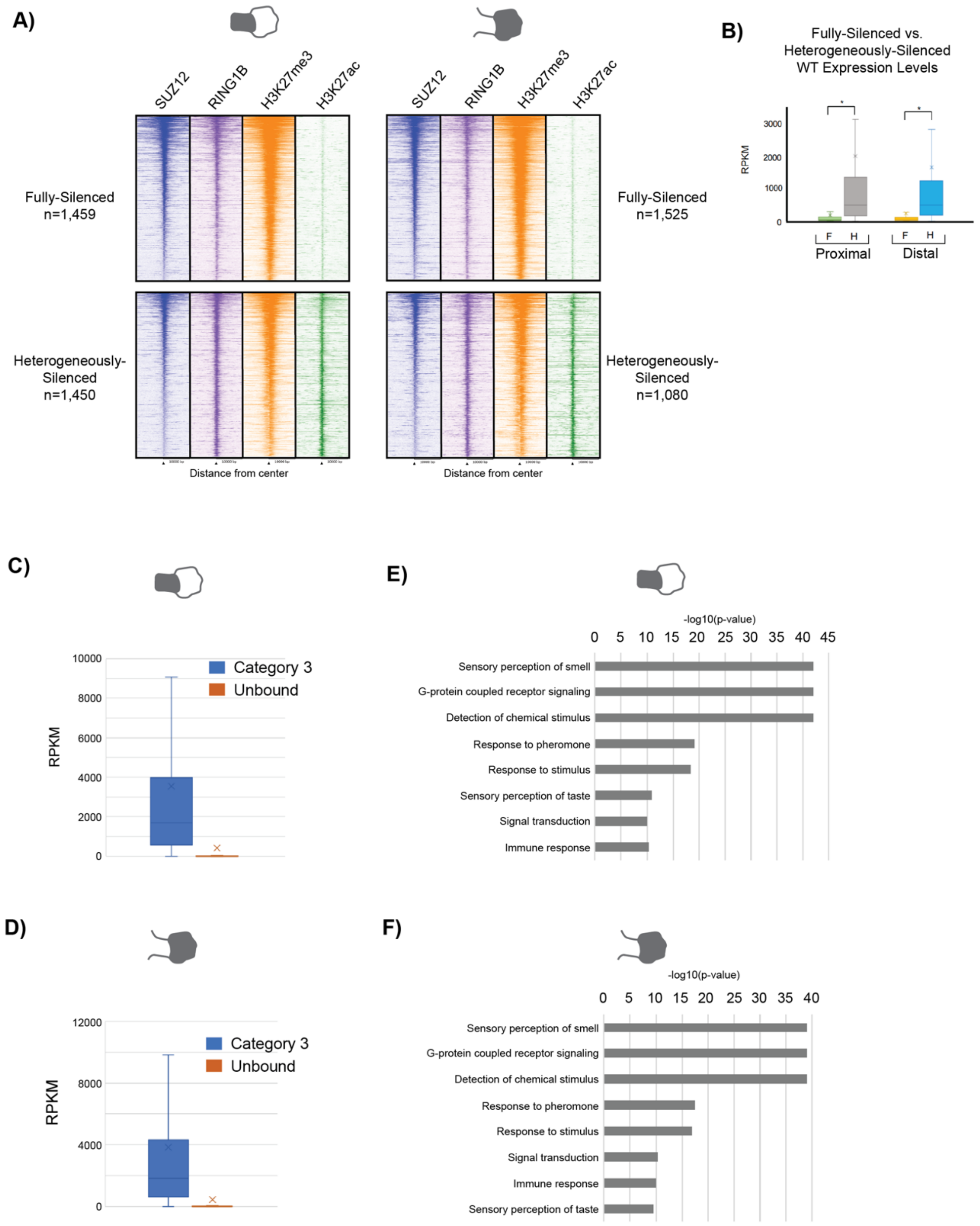


**Figure S1. PRC1 and PRC2 co-occupancy is sub-categorized to account for cell heterogeneity** **(A)** Heatmaps of ChIP-seq for SUZ12 (blue), RING1B (purple), H3K27me3 (orange), and H3K27ac (green) at promoters that are fully-silenced or heterogeneously-silenced as distinguished

by absence or presence of H3K27ac, respectively. The number (n) of promoters that fall into either sub-category are indicated.
**(B)** Box plot of gene expression (RPKM values) comparing fully-silenced genes (indicated by 'F') and heterogeneously-silenced genes (indicated by 'H') in wild type proximal and distal limbs.
**(C-D)** Box plot of gene expression (RPKM values) comparing unbound promoters and category 3 (RING1B bound) promoters in proximal and distal limb, respectively.
**(E-F)** Gene Ontology terms (using DAVID) of top 8 biological processes associated with genes that are unbound by PRC in proximal and distal limb, respectively.

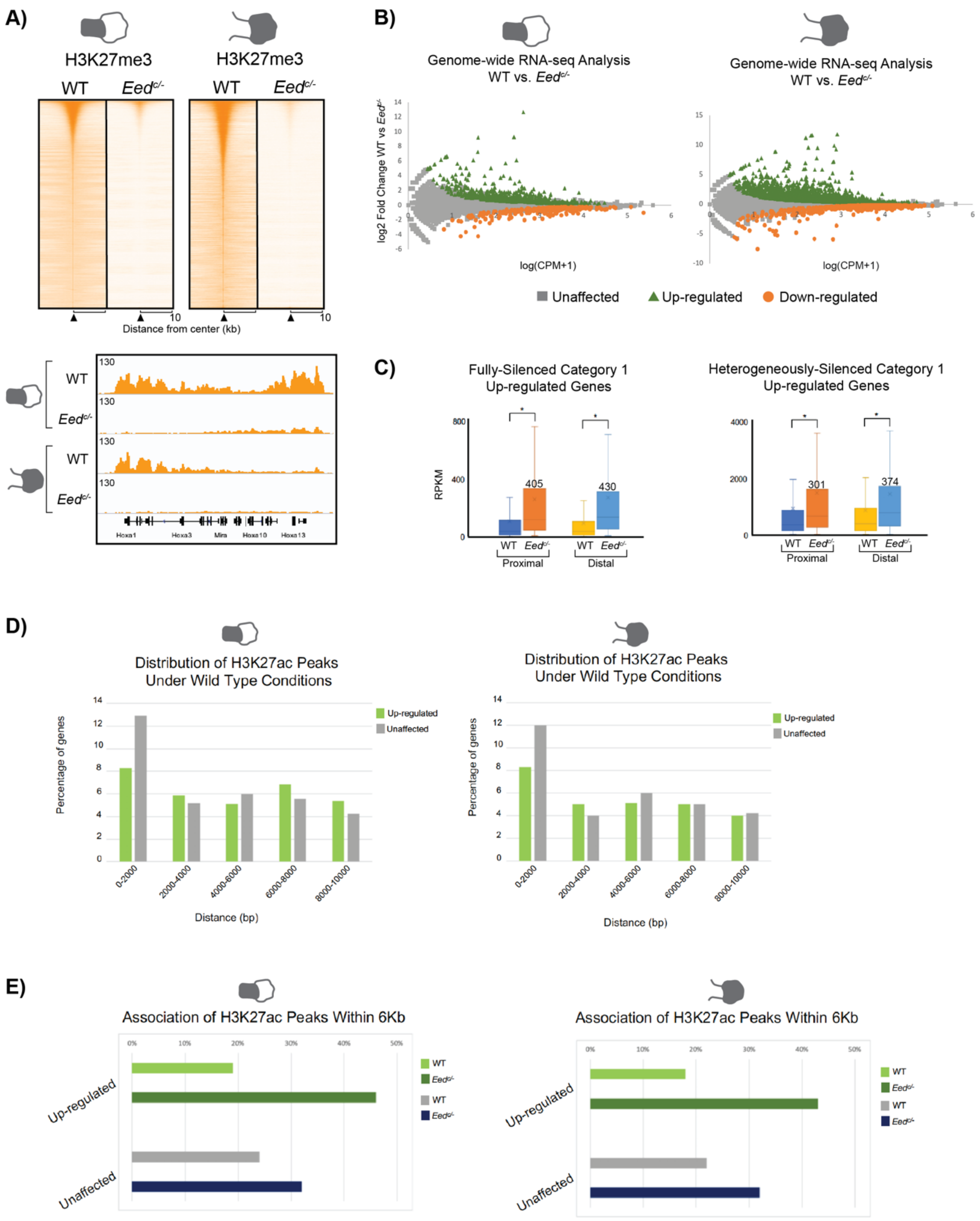


**Figure S2. PRC2 inactivation results in the up-regulation of fully- and heterogeneously-silenced genes**

**(A)** Heatmaps of ChIP-seq for genome-wide H3K27me3 in wild type and *Eed*$^{c/-}$ proximal and distal limb tissue (upper panel). IGV snapshot of H3K27me3 over the *HoxA* gene cluster show a reduction in H3K27me3 levels in *Eed*$^{c/-}$ proximal and distal limbs.
**(B)** MA plots depicted genome-wide gene expression changes upon *Eed* inactivation in proximal and distal limb. Unaffected genes are depicted in grey, up-regulated genes in green and down-regulated genes in orange.
**(C)** Box plots of expression levels (RPKM values) of genes that are up-regulated upon *Eed* inactivation from the categories of promoters that are both fully- and heterogeneously-silenced in wild type and *Eed*$^{c/-}$ proximal and distal limbs. The number of up-regulated genes is indicated on the bar graph.
**(D)** Bar graphs representing the distribution of H3K27ac peaks to promoters of genes that are either up-regulated or unaffected by *Eed* inactivation under wild type conditions in proximal and distal limbs, according to distance.
**(E)** Bar graphs representing the percentage of up-regulated and unaffected genes that are associated to a H3K27ac peak within a 6kb distance of the gene promoter in wild type and *Eed*$^{c/-}$ proximal and distal limbs.

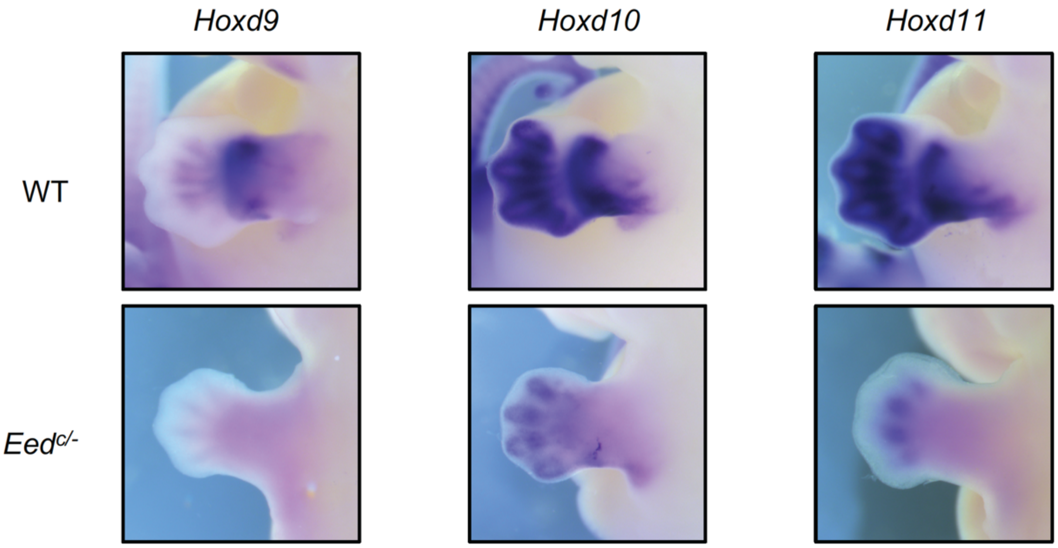


**Figure S3. Impaired regulation of the *Hoxd9*, *10*, and *11* genes upon PRC2 inactivation**
Whole-mount *in situ* hybridizations of *Hoxd9, Hoxd10* and *Hoxd11* genes in wild type and *Eed*$^{c/-}$ limb buds.

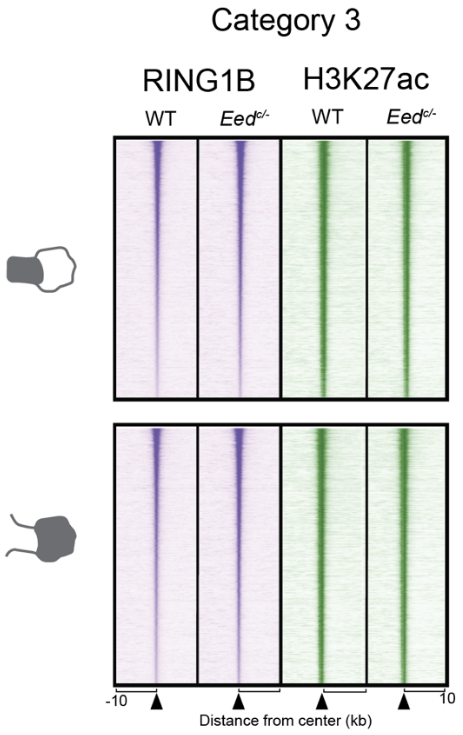


**Figure S4. RING1B occupancy at category 3 promoters is unaffected by PRC2 inactivation**
Heatmaps of ChIP-seq signals for RING1B (purple) and H3K27ac (green) in wild type and *Eed*$^{c/-}$ proximal and distal limbs at promoters of genes exclusively bound by PRC1 under wild type conditions.
